# Is asthma associated with exposure to smoke from a coal mine fire?

**DOI:** 10.1101/631317

**Authors:** S Taylor, B Borg, C Gao, D Brown, R Hoy, A Makar, T McCrabb, J Ikin, BR Thompson, MJ Abramson

## Abstract

In 2014, a fire at the Hazelwood open cut coal mine (Victoria, Australia) burned for about 6 weeks. Residents of the adjacent town of Morwell were exposed to high levels of fine particulate matter (PM_2.5_) during this period. Three and a half years after the event, this study aimed to investigate the long-term impact of short-term exposure to coal mine fire smoke on asthma.

A cross-sectional analysis was undertaken on a group of exposed participants with asthma from Morwell (n=165) and a group of unexposed participants with asthma from the control town of Sale (n=64). Town exposure status was determined by modelled PM_2.5_ data for the mine fire period. Respiratory symptoms were assessed with a validated respiratory health questionnaire and symptom severity score. Asthma control was assessed with an asthma control questionnaire. Lung function testing included spirometry, bronchodilator response, and fraction of exhaled nitric oxide.

There was no evidence that exposed Morwell participants had more severe asthma symptoms, worse lung function, or more eosinophilic airway inflammation compared to unexposed Sale participants. However there was some evidence that Morwell participants had more uncontrolled than well-controlled asthma, compared to the participants from Sale (adjusted relative risk ratio 2.71 95%CI: 1.02, 7.21, p=0.046).

Three and a half years after exposure, coal mine fire smoke does not appear to be associated with more severe asthma symptoms or worse lung function, but may be associated with poorer asthma control.

**Summary take home message:** In people with asthma, short-term coal mine fire smoke exposure does not appear to have long-term impact on severity of asthma symptoms, lung function or eosinophilic airway inflammation, but may affect asthma control.

## Introduction

Air pollution, from either natural or anthropogenic sources, such as forest fires, motor vehicles and industry emissions, is a well-recognised environmental trigger of asthma[1-3]. Exposure to airborne particulate matter (PM) is known to cause exacerbations of asthma, impair lung function, and possibly contribute to the development of asthma[1, 3-10]. Of particular concern is exposure to respirable particles with a median aerodynamic diameter less than 10μm (PM_10_). Furthermore, fine particles less than 2.5um (PM_2.5_) have been of greater interest as these fine particles move to the periphery of the lung, and there is evidence they are associated with greater respiratory effects, including more severe asthma symptoms[3, 7, 8, 11].

In February 2014, a fire started in the Hazelwood open cut brown coal mine, located in the Latrobe Valley, Victoria, Australia. It was an unprecedented event that generated significant air pollution from coal mine fire smoke over a six week period, predominantly affecting residents in the adjacent town of Morwell (population approximately 14 000[12]). This resulted in considerable community concern about the potential long-term health effects of the smoke exposure, much of which related to the respiratory health impacts.

To the best of our knowledge, there was no evidence in the literature about the impact of short-term coal mine fire smoke exposure on asthma[5]. However, there was some evidence for the impact of analogous exposures on respiratory health. For example, exposure to PM_2.5_ and PM_10_ from wildfire smoke has been associated with a concomitant increase in severity of asthma symptoms, asthma medication use, asthma-related emergency presentations and hospital admissions[13-19]. Yet there were no studies that looked at the long-term health impacts of wildfire smoke in the years following these events. Another example was domestic coal combustion, smoke from which is of similar composition to coal mine fire smoke. This was associated with increased asthma symptoms, childhood asthma and reduced lung function[20-23]. However this smoke exposure tended to be over a period of years, and it is not known if the respiratory impacts are similar for a one-off exposure event.

The Hazelwood Health Study (HHS; www.hazelwoodhealthstudy.org.au) was established to investigate long-term health outcomes in people who were exposed to smoke from the Hazelwood mine fire. Findings from the 2016 Adult Survey demonstrated that, compared to a control population from the unexposed Victorian town of Sale[24], the exposed population of Morwell had a significant increase in self-reported asthma symptoms, asthma diagnoses since 2014, and a higher mean symptom severity score amongst asthmatics[25].

This analysis aimed to further investigate health findings in relation to asthma. Specifically, three and a half years after the fire, we examined whether a group of people with asthma exposed to the coal mine fire smoke had more severe asthma symptoms, worse lung function and poorer control, compared to a group of unexposed people with asthma.

## Methods

### Study design and setting

This cross-sectional analysis draws upon data from round one of respiratory testing in the longitudinal HHS Respiratory Stream. Residents of Morwell and Sale had previously been selected as the ‘exposed’ and ‘unexposed’ populations based upon mine fire-related hourly PM_2.5_ exposure levels modelled by The Commonwealth Scientific and Industrial Research Organisation (CSIRO) Oceans & Atmosphere Flagship (see fig. 1)[24]. During the mine fire period, the high-resolution modelling showed that PM_2.5_ levels exceeded Australian air quality standards (25µg m^-3^ as a 24-hour average) on 23 days in southern Morwell and 12 days in eastern Morwell[24]. In the first two days of the fire, PM_2.5_ concentrations were predicted to have reached up to 3700µg m^-3^ in southern Morwell[24]. There were no air quality breaches in Sale, a regional Victorian town that had selected areas with comparable socio-economic status, age distribution, household size and population stability as Morwell[25]. Therefore Sale provided an appropriate unexposed control population. Clinical testing for the Respiratory Stream was conducted in Morwell between August and December 2017, and in Sale between January and March 2018.

**Figure 1.**
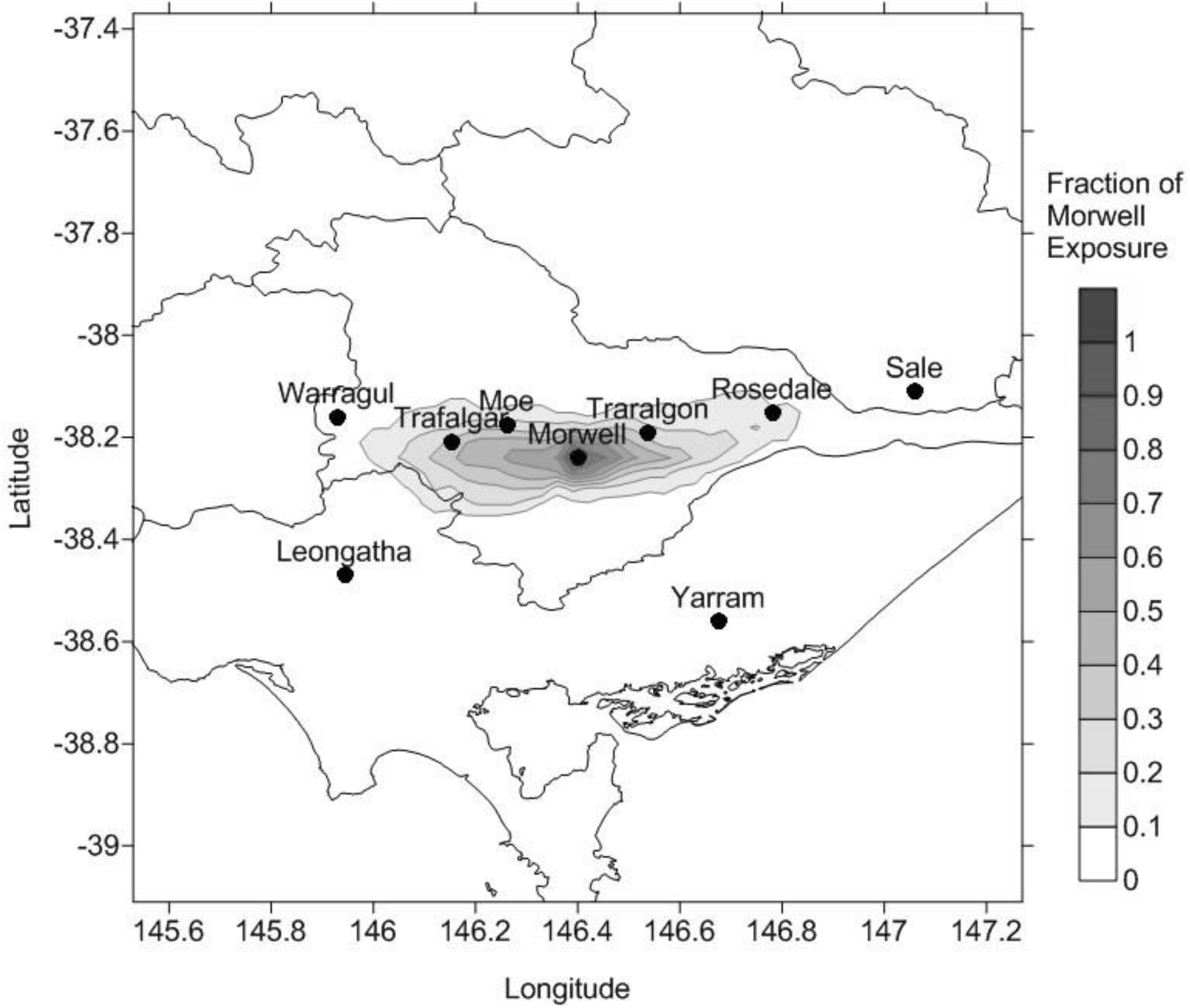
Modelled smoke exposure during the Hazelwood mine fire, for towns in the Latrobe Valley including Morwell and Sale. *Source:* Emmerson K, Reisen R, Luhar A, Williamson G, Cope M. Air Quality Modelling of Smoke Exposure from the Hazelwood Mine Fire. CSIRO Australia, 2016. www.hazelwoodhealthstudy.org.au/study-findings/study-reports/ Date last updated: December 2016. Date last accessed: 21 January 2019.

### Participants

Potential participants for the Respiratory Stream were drawn from the Hazelwood Adult Survey cohort. Methodology for the Adult Survey has been outlined previously[25]. To summarise briefly, Adult Survey participants must have been aged at least 18 years and have lived in Morwell or in selected areas of Sale at the time of the mine fire. Adult Survey participants were excluded from eligibility for the Respiratory Stream if they had requested no further contact after the Adult Survey, had unknown age or sex, or were aged over 90 years.

From 3,854 eligible Adult Survey participants, a weighted random sample of 1,346 people was selected to be invited in to the Respiratory Stream. Adult Survey participants who had reported having asthma were intentionally over sampled such that they represented 40% of those invited from both Morwell and Sale. For the first round of testing, the desired number of Respiratory Stream participants was 339 in Morwell and 170 in Sale. This sample size calculation assumed a mean FEV1 decline in the non-exposed community of 23.1 (±17.1) ml/year[39], based on a two-sample t-test, a significance level of p<0.05 and power of 0.90. The numbers of asthmatics expected to participate were approximately 136 in Morwell and 68 in Sale.

Recruitment was via mailed invitation, followed by reminder mail and phone calls to non-responders. Upon contact, potential participants had to be excluded if they had a contraindication to spirometry, such as recent surgery, myocardial infarction, pneumothorax, pulmonary embolus, open pulmonary tuberculosis or known aneurysms[26].

### Demographics

For each participant, demographic characteristics, such as age, sex, ethnicity, employment status, educational background and occupational exposure (working in dusty or polluted environments, such as coal mines, farms or driving diesel trucks) had been previously collected by self-report at the time of the Adult Survey[25].

### Medication use and health risk factors

All data collection in the Respiratory Stream clinic was undertaken by trained respiratory scientists. Study data were managed using REDCap (Research Electronic Data Capture)[27] electronic data capture tools hosted at Monash University (Victoria, Australia).

An interviewer-administered, standardised questionnaire recorded asthma medication use in the previous three months, including type and dosage regimen[28]. Corticosteroid doses were calculated as beclomethasone daily dose equivalents (mcg), which were then categorised into low (100-200mcg), medium (>200-400mcg) or high (>400mcg) [2]. Smoking history was taken and participants were classified as current smokers, former smokers (current non-smokers with >100 cigarettes in their lifetime) or never smoked (< 100 cigarettes in their lifetime)[29]. Duration of smoking (years) was also calculated for current and former smokers[30]. A self-reported history of nasal allergies or hay fever was recorded as an indicator of atopy. Physical characteristics, including height and weight, were measured and body mass index (BMI) was calculated[31].

### Self-perceived respiratory health

Self-perceived respiratory symptoms and conditions were assessed via an interviewer-administered questionnaire adapted from the second European Community Respiratory Health Study (ECRHSII)[28]. Diagnosis of asthma was by self-report. An asthma symptom severity score, designed by Pekkanan et al.[32], was calculated from eight questions included in the questionnaire. Asthma control was assessed with the validated, self-administered, 7-item Asthma Control Questionnaire (ACQ-7)[33]. Responses were given on a 7-point scale, with the score being a mean of the responses. Recognised cut points were used to categorise scores into having well-controlled asthma (ACQ < 0.75), borderline control (0.75 ≥ ACQ ≤ 1.5), and uncontrolled asthma (ACQ > 1.5)[34].

### Respiratory function tests

Standardisation between staff and testing sites was achieved through the use of Standard Operating Procedures and validated equipment. Preparation instructions were provided to participants at the time of booking the tests. Medication was not withheld prior to respiratory testing for ethical reasons. Instead, the time inhaled medication was last taken was recorded at testing, and this was adjusted for at the analysis stage.

Respiratory function was determined with spirometry using the EasyOne Pro™ LAB Respiratory Analysis System (nnd Medical Technologies AG, Zurich, Switzerland) performed according to American Thoracic Society/European Respiratory Society (ATS/ERS) criteria[35]. Parameters measured included forced expiratory volume in 1 s (FEV_1_), forced vital capacity (FVC) and FEV_1_/FVC. Z-scores for FEV_1_, FVC and FEV_1_/FVC were calculated using the Global Lung Initiative 2012 spirometry reference equations[36]. The measurements were repeated ten minutes after administration of three puffs of Salbutamol pressurised Metered Dose Inhaler (pMDI)(100ug per activation)[37]. Mean percentage change in FEV_1_ for each exposure group was calculated.

Fraction of exhaled nitric oxide (FeNO) was measured using the Niox Vero FeNO device (Aerocrine, Solna, Sweden) according to ATS/ERS recommendations[38]. A mean FeNO was calculated for each group.

### Statistical methods

Statistical analysis and data transformations were performed using Stata version 15 (Stata Corporation, College Station, Texas 2017). Simple descriptive statistics were first used to summarise characteristics and clinical outcomes for the exposed (Morwell) and non-exposed (Sale) participants, with crude differences between the two samples assessed using Pearson chi-squared tests for categorical measures and t-tests for continuous measures. Linear regression models were used to compare the univariate and multivariate differences (controlling for key confounders) in continuous outcomes between the exposed and non-exposed samples. FeNO was transformed to log scale in linear regression due to its skewed distribution. Logistic regression models were used for binary outcomes, and multinomial logistic regression models were used for the categorical outcomes. Multiple-imputation (MI) procedures using chained equation were used in regression models to obtain more accurate estimates, and to control for non-response bias.

### Ethical considerations

The Monash University Human Research Ethics Committee (MUHREC) approved the Hazelwood Health Study: Cardiovascular and Respiratory Streams (approval number 1078). All participants provided written informed consent.

## Results

As shown in Figure 2, the Respiratory Stream recruited 346 Morwell participants and 173 Sale participants, totalling 519 (39% of those eligible). For this cross-sectional analysis, only data from the 229 asthmatic participants were analysed; 165 from Morwell and 64 from Sale.

**Figure 2.**
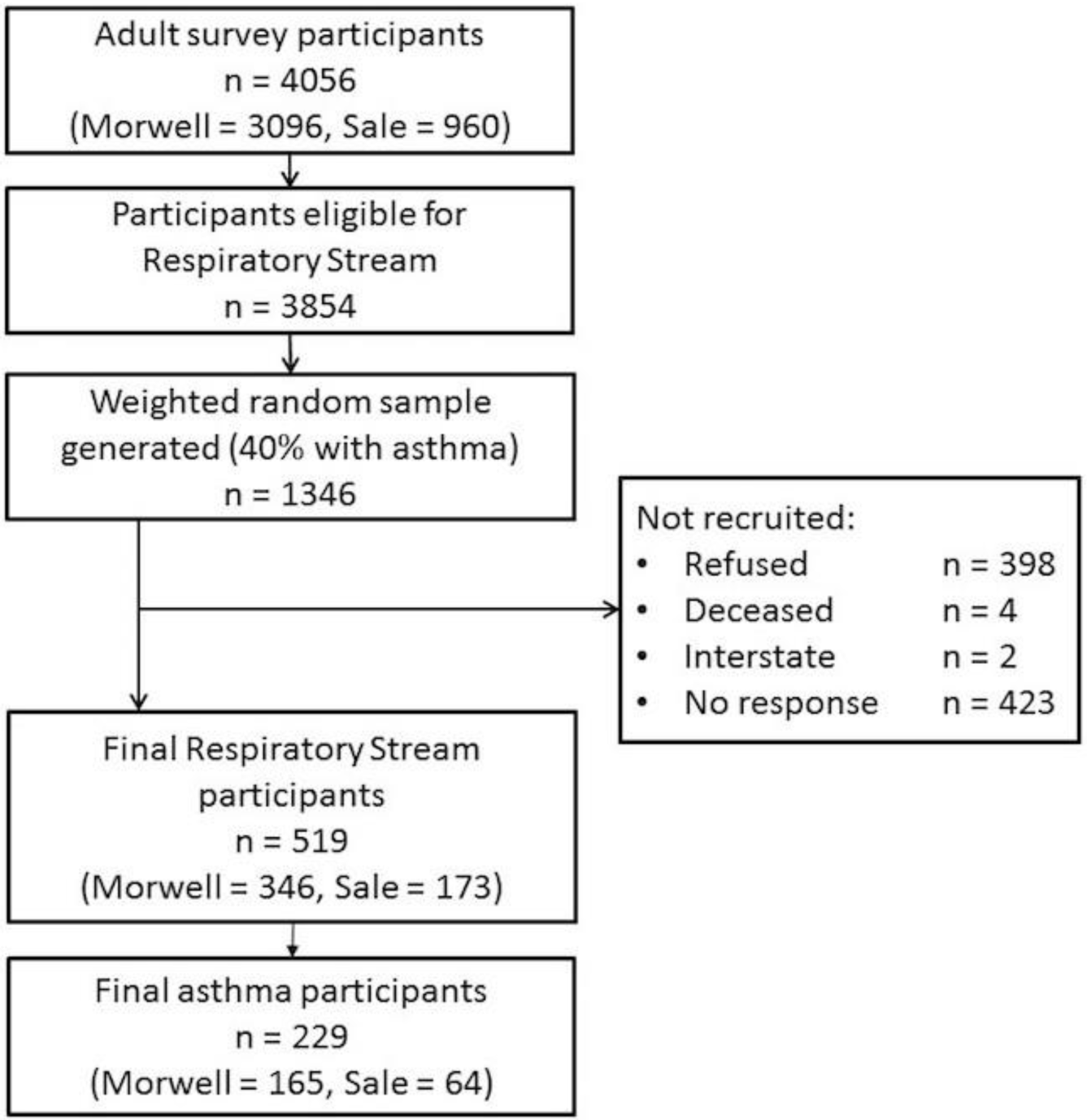
Participant recruitment flow diagram.

The demographic characteristics of the participants from Morwell and Sale are shown in Table 1. Participants from Morwell were more likely to be using medium to high doses of inhaled corticosteroids. Otherwise Morwell and Sale participants were similar.

**Table 1.** Demographic characteristics for participants from Morwell and Sale.

| Participant characteristics | Morwell<br>N=165 |  | Sale<br>N=64 |  | p-value |
| --- | --- | --- | --- | --- | --- |
|  | n | % | n | % |  |
| **Age/years** |  |  |  |  | 0.376 |
| 18-29 | 13 | 7.9 | 3 | 4.7 |  |
| 30-44 | 45 | 27.3 | 13 | 20.3 |  |
| 45-59 | 52 | 31.5 | 20 | 31.3 |  |
| 60-74 | 39 | 23.6 | 23 | 35.9 |  |
| 75+ | 16 | 9.7 | 5 | 7.8 |  |
| **Gender** |  |  |  |  | 0.948 |
| Male | 56 | 33.9 | 22 | 34.4 |  |
| Female | 109 | 66.1 | 42 | 65.6 |  |
| **Ethnicity** |  |  |  |  | 0.532 |
| Caucasian / White | 164 | 99.4 | 64 | 100 |  |
| Other | 1 | 0.6 | 0 | 0 |  |
| **Employment status** |  |  |  |  | 0.320 |
| Employed | 64 | 38.8 | 31 | 48.4 |  |
| Retired | 44 | 26.7 | 19 | 29.7 |  |
| Not working due to poor health | 20 | 12.1 | 6 | 9.4 |  |
| Other (unemployed, studying, home duties etc.) | 37 | 22.4 | 8 | 12.5 |  |
| **Highest educational qualification** |  |  |  |  | 0.195 |
| Secondary up to year 10 | 40 | 24.5 | 13 | 20.6 |  |
| Secondary year 11-12 | 34 | 20.9 | 10 | 15.9 |  |
| Certificate (trade/apprenticeship/technicians) | 53 | 32.5 | 30 | 47.6 |  |
| University or other Tertiary Institute degree | 36 | 22.1 | 10 | 15.9 |  |
| **Body Mass Index (BMI)** |  |  |  |  | 0.671 |
| Underweight/Normal (BMI<25kg/m<sup>2</sup>) | 27 | 16.4 | 11 | 17.2 |  |
| Overweight (25≤BMI<30) | 43 | 26.1 | 20 | 31.3 |  |
| Obese (BMI≥30) | 95 | 57.6 | 33 | 51.6 |  |
| **Smoking status** |  |  |  |  | 0.129 |
| Never | 80 | 48.5 | 22 | 34.4 |  |
| Former smoker | 50 | 30.3 | 28 | 43.8 |  |
| Current smoker | 35 | 21.2 | 14 | 21.9 |  |
| **Nasal allergies/Hay fever** |  |  |  |  | 0.105 |
| Yes | 102 | 61.8 | 32 | 50 |  |
| No | 63 | 38.2 | 32 | 50 |  |
| **Asthma medication** |  |  |  |  | 0.834 |
| No inhaled medications | 66 | 40 | 26 | 40.6 |  |
| Other asthma medications* | 34 | 20.6 | 11 | 17.2 |  |
| Inhaled corticosteroids | 65 | 39.4 | 27 | 42.2 |  |
| **Inhaled corticosteroid use<sup>†</sup>** |  |  |  |  | 0.001 |
| No use | 101 | 61.2 | 40 | 62.5 |  |
| Low dose (100-200mcg) | 16 | 9.7 | 17 | 26.6 |  |
| Medium dose (>200-400mcg) | 21 | 12.7 | 2 | 3.1 |  |
| High dose (>400mcg) | 27 | 16.4 | 5 | 7.8 |  |
| **Occupational exposure<sup>‡</sup>** |  |  |  |  |  |
| Exposed | 57 | 34.5 | 26 | 40.6 | 0.394 |
|  | **Mean** | **SD** | **Mean** | **SD** | **p-value** |
| **Years of smoking among current and former smokers (n=127)** | 20.7 | 12.5 | 24.2 | 13.8 | 0.212 |
| **Age** | 52.1 | 16.3 | 55.7 | 14.8 | 0.108 |

\*Other asthma medications including short-acting beta agonists, short-acting muscarinic antagonists, long-acting beta agonists (LABA), long-acting muscarinic antagonists (LAMA), LAMA/LABA combinations.
<sup>†</sup>Doses were calculated as beclomethosone equivalents (mcg).
<sup>‡</sup>Occupational exposure from working in dusty or polluted environments, such as coal mines, farms and driving diesel trucks.

The primary outcomes for Morwell and Sale are reported in Table 2. For all the self-reported asthma symptom questions, a higher proportion of Morwell participants reported symptoms compared with Sale. However, there was no evidence that the unadjusted symptom proportions differed statistically between the two samples. Other unadjusted outcomes are comparable between the groups except for a slightly higher proportion of uncontrolled asthma in Morwell.

**Table 2.** Respiratory questionnaires and lung function outcomes for participants from Morwell and Sale.

| Outcome | Morwell<br>N=165 |  | Sale<br>N=64 |  | p-value |
| --- | --- | --- | --- | --- | --- |
|  | n | % | n | % |  |
| **First asthma attack post 2014** | 18 | 11 | 8 | 12.9 | 0.688 |
| **Most recent asthma attack post 2014** | 117 | 70.9 | 41 | 66.1 | 0.478 |
| **Asthma-related respiratory symptoms** |  |  |  |  |  |
| Wheeze in the last 12 months | 113 | 68.5 | 43 | 67.2 | 0.852 |
| Wheeze and breathlessness | 90 | 54.5 | 33 | 51.6 | 0.691 |
| Wheeze without URTI | 94 | 57 | 35 | 54.7 | 0.754 |
| Chest tightness in last 12 months | 61 | 37 | 15 | 23.4 | 0.057 |
| Dyspnoea at rest in last 12 months | 47 | 28.5 | 15 | 23.4 | 0.427 |
| Dyspnoea after exercise in last 12 months | 108 | 65.9 | 35 | 54.7 | 0.121 |
| Woken with dyspnoea in last 12 months | 37 | 22.4 | 9 | 14.1 | 0.162 |
| Woken with cough in last 12 months | 91 | 55.2 | 28 | 43.8 | 0.145 |
| Asthma attack in last 12 months | 76 | 46.1 | 28 | 43.8 | 0.754 |
|  | **Mean** | **SD** | **Mean** | **SD** | **p-value** |
| **Spirometry\*** |  |  |  |  |  |
| FEV<sub>1</sub> L | 2.7 | 0.8 | 2.5 | 0.7 | 0.164 |
| FEV<sub>1</sub> (z-score) | -0.8 | 1.1 | -0.9 | 1.2 | 0.665 |
| FEV<sub>1</sub> % Predicted | 88.5 | 16.8 | 86.2 | 18.4 | 0.465 |
| FVC L | 3.5 | 1.0 | 3.5 | 0.8 | 0.616 |
| FVC (z-score) | -0.4 | 1.0 | -0.3 | 1.0 | 0.514 |
| FVC % Predicted | 94.8 | 14.4 | 96.0 | 14.9 | 0.609 |
| FEV<sub>1</sub>/FVC | 74.6 | 10.1 | 71.3 | 11.7 | 0.072 |
| FEV<sub>1</sub>/FVC (z-score) | -0.7 | 1.1 | -1.0 | 1.2 | 0.141 |
| FEV<sub>1</sub>/FVC % Predicted | 92.8 | 11.8 | 89.4 | 13.2 | 0.100 |
| **Post BD response\*** |  |  |  |  |  |
| % change FEV<sub>1</sub> | 6.4 | 7.2 | 8.0 | 8.0 | 0.154 |
| **FeNO\*** | 21.4 | 17.8 | 27.0 | 22.2 | 0.066 |
| **Asthma symptom severity score** | 4.1 | 2.2 | 3.7 | 2.0 | 0.140 |
|  | **n** | **%** | **n** | **%** | **p-value** |
| **Asthma control (ACQ-7)** |  |  |  |  | 0.255 |
| Well-controlled asthma (ACQ < 0.75) | 90 | 56.3 | 42 | 65.6 |  |
| Borderline controlled asthma (ACQ 0.75-1.5) | 35 | 21.9 | 14 | 21.9 |  |
| Uncontrolled asthma (ACQ >1.5) | 35 | 21.9 | 8 | 12.5 |  |

URTI: upper respiratory tract infection; FEV<sub>1</sub>: forced expiratory volume in 1 s; FVC: forced vital capacity; BD: bronchodilator; FeNO: fraction of exhaled nitric oxide; ACQ-7: 7-item asthma control questionnaire[33].
\*Results were missing for five spirometry, nine post BD response, and one FeNO test result due to difficulty in conducting the test or low test quality.

Table 3 and Table 4 show univariate and multivariate regression analyses investigating the association between exposure (Morwell versus Sale) and a selection of the primary outcomes. After adjustment for confounders, there was no association with Morwell or Sale for any of the self-reported asthma symptoms or objective measures of lung function and airway inflammation.

**Table 3.** Linear regression analysis of the association between exposure and primary outcomes.

|  | Univariate regression<br>Mean diff<br>(95% CI) | Multivariate regression<br>Adj mean diff<br>(95% CI) | p-value |
| --- | --- | --- | --- |
| FEV<sub>1</sub> (z-score) | 0.07 (-0.31,0.46) | 0.06 (-0.29,0.40)* | 0.741 |
| FEV<sub>1</sub>/FVC (z-score) | 0.28 (-0.11,0.67) | 0.25 (-0.12,0.62)* | 0.180 |
| % change FEV<sub>1</sub> (BD response) | -1.66 (-3.98,0.67) | -1.50 (-3.72,0.73)<sup>†</sup> | 0.186 |
| Log FeNO | -0.18 (-0.40,0.04) | -0.20 (-0.42,0.03)<sup>‡</sup> | 0.084 |
| Asthma symptom severity score | 0.44 (-0.14,1.02) | 0.31 (-0.20,0.81)<sup>§</sup> | 0.229 |

\*Adjusted for employment status, education, smoking status, years of smoking, occupational exposure, nasal allergies/hay fever, and whether the patient took asthma medication prior to lung function testing.
<sup>†</sup>Adjusted for age, gender, body mass index, height, employment status, education, smoking status, years of smoking, occupational exposure, nasal allergies/hay fever, and whether the patient took asthma medication prior to lung function testing.
<sup>‡</sup>Adjusted for age, gender, body mass index, height, employment status, education, current smoking, inhaled corticosteroid dose, nasal allergies/hay fever, and whether the patient followed the FeNO preparation guidelines.
<sup>§</sup>Adjusted for age, gender, body mass index, employment status, education, smoking status, occupational exposure, nasal allergies/hay fever, and asthma medication use.
FEV<sub>1</sub>: forced expiratory volume in 1 s; FVC: forced vital capacity; BD: bronchodilator; FeNO: fraction of exhaled nitric oxide.

**Table 4.** Logistic regression analysis of the association between exposure and primary outcomes.

|  | Univariate regression<br>Crude OR<br>(95% CI) | Multivariate regression<br>Adj OR*<br>(95% CI) | p-value |
| --- | --- | --- | --- |
| Wheeze in the last 12 months | 1.06 (0.57,1.99) | 0.90 (0.41,1.97) | 0.798 |
| Wheeze and breathlessness | 1.13 (0.62,2.04) | 0.95 (0.46,1.96) | 0.886 |
| Wheeze without URTI | 1.10 (0.61,1.96) | 1.07 (0.55,2.09) | 0.838 |
| Chest tightness in last 12 months | 1.92 (0.98,3.76) | 2.11 (0.93,4.77) | 0.073 |
| Dyspnoea at rest in last 12 months | 1.30 (0.68,2.50) | 1.04 (0.47,2.29) | 0.920 |
| Dyspnoea after exercise in last 12 months | 1.60 (0.88,2.90) | 1.66 (0.83,3.30) | 0.148 |
| Woken with dyspnoea in last 12 months | 1.77 (0.79,3.96) | 1.77 (0.71,4.41) | 0.217 |
| Woken with cough in last 12 months | 1.58 (0.85,2.94) | 1.66 (0.84,3.25) | 0.141 |
| Asthma attack in last 12 months | 1.10 (0.61,1.98) | 1.05 (0.49,2.26) | 0.897 |

\*Adjusted for age, gender, body mass index, education, employment status, smoking status, occupational exposure, nasal allergies/hay fever, and asthma medication use.
URT: upper respiratory tract infection.

When the ACQ-7 results were compared (Table 5), there was no effect for exposure (Morwell versus Sale) when those with borderline control (0.75 ≥ ACQ ≤ 1.5) were compared to those who were well-controlled (ACQ < 0.75). However, after adjustment for confounders, compared to Sale participants, Morwell participants had comparatively more uncontrolled asthma (ACQ > 1.5) than well-controlled asthma (p=0.046). Being a current smoker and being on any type of asthma medication was positively associated with being borderline or uncontrolled compared to being well-controlled.

**Table 5.** Multinomial logistic regression analysis of borderline versus well-controlled asthma and uncontrolled versus well-controlled asthma for ACQ-7*.

|  | Borderline vs well-controlled |  |  | Uncontrolled vs well-controlled |  |  |
| --- | --- | --- | --- | --- | --- | --- |
|  | Crude RRR | Adj RRR<sup>†</sup> (95% CI) | p-value | Crude RRR | Adj RRR<sup>†</sup> (95% CI) | p-value |
| **Morwell versus Sale** | 1.16 | 1.28 (0.53, 3.13) | 0.585 | 2.25 | 2.71 (1.02, 7.21) | **0.046** |
| **Smoking status** |  |  |  |  |  |  |
| Never | Ref | Ref |  | Ref | Ref |  |
| Former smoker | 2.05 | 1.75 (0.73, 4.22) | 0.209 | 0.84 | 0.58 (0.21, 1.59) | 0.288 |
| Current smoker | 2.59 | 5.82 (1.74, 19.49) | **0.004** | 2.00 | 3.70 (1.19, 11.53) | **0.024** |
| **Asthma medication** |  |  |  |  |  |  |
| No | Ref | Ref |  | Ref | Ref |  |
| Other asthma medications | 4.92 | 5.66 (1.69, 18.91) | **0.005** | 8.91 | 9.75 (3.11, 30.56) | **<0.001** |
| Inhaled corticosteroids | 7.10 | 9.08 (3.20, 25.79) | **<0.001** | 13.76 | 17.44 (5.81, 52.38) | **<0.001** |

\*ACQ-7: 7-item asthma control questionnaire, where well-controlled asthma ( $\text{ACQ} < 0.75$ ), borderline controlled asthma ( $\text{ACQ} 0.75\text{--}1.5$ ) and uncontrolled asthma ( $\text{ACQ} > 1.5$ ).
<sup>†</sup>Adjusted for age, gender, BMI, employment status, education, smoking status, asthma medication use, and nasal allergies/hay fever.
RRR: relative risk ratio; BMI: body mass index.

## Discussion

Nearly four years after the Hazelwood mine fire, there was no evidence that exposed participants have significantly more severe asthma symptoms, worse lung function, or more eosinophilic airway inflammation compared to unexposed participants, as assessed by the asthma symptom questionnaire, severity score, spirometry or FeNO. However, compared to the participants from Sale, there was evidence that participants from Morwell have more uncontrolled than well-controlled asthma when assessed with the validated ACQ-7 (Table 5).

One possible reason for our finding of difference in asthma control is that particulates in the mine fire smoke caused long-lasting respiratory damage, potentially reducing the ability of some exposed participants to achieve asthma control in the years after the fire. However, prior to the HHS, there had been no published evidence relating to the long-term respiratory health impacts of short-term coal mine fire smoke exposure[5]. There is evidence from ambient air pollution studies that acute exposure to PM causes damage to the respiratory system through oxidative stress and inflammation, impacting on respiratory function and resulting in exacerbations of asthma[3, 6, 39]. Furthermore, long-term exposure to PM from ambient pollution is known to cause chronic inflammation and airway remodelling, resulting in ongoing respiratory morbidity[3, 6, 39]. However, it is still unknown whether short-term exposure to PM from coal mine fire smoke causes long-term respiratory damage by similar pathophysiological mechanisms, if at all. Arguably, the lack of positive results for the other respiratory outcomes we examined makes it less likely that chronic respiratory damage from the smoke exposure has played a role in the asthma control findings.

Additionally, because the ACQ only assessed asthma control in the week prior to completing the questionnaire, other factors in Morwell at the time of testing may have been responsible for the difference in asthma control findings. Testing of Morwell subjects coincided with the grass pollen season in Victoria (October to December). We attempted to adjust for pollen season by taking into account nasal allergies and hay fever, however it is possible that this still had an impact on control. We also could not exclude the role of other unknown environmental factors during the testing period.

The finding of no significant difference between exposed and unexposed samples for self-reported asthma symptoms and the asthma symptom severity score was inconsistent with the results of the larger Adult Survey, which found exposed participants had significantly increased asthma-related respiratory symptoms, and a significantly higher mean modified asthma symptom severity score[25]. The Adult Survey was a population-based study that examined the prevalence of respiratory symptoms in the general population, and included not only people with asthma, but also those with chronic obstructive pulmonary disease and other respiratory conditions. A possible reason for the inconsistency between the Adult Survey and this sub-study is that a much larger number of people were surveyed in the Adult Survey, including a greater number of asthmatics overall, giving it greater power to detect differences in respiratory findings. However, our results may also mean that on further examination, there was no significant difference in the severity of asthma symptoms between people with asthma from Morwell and Sale.

Interestingly, we found a higher proportion of Morwell residents were on medium to high dose inhaled corticosteroids. It is possible that the Morwell participants had been prescribed higher doses of inhaled corticosteroids because they had more severe asthma, potentially due to mine fire smoke exposure. We may not have seen a difference in the self-reported asthma symptoms, symptom severity score or lung function results in this analysis because the participants had been adherent with their steroid medication.

### Strengths and limitations

The main strength of this analysis is that it provides evidence that suggests short-term coal mine fire smoke exposure has limited long-term impact on asthma. To the best of our knowledge, there are no other similar studies in the literature examining this issue, so our study adds to the evidence base. A further strength is that we used validated questionnaires and objective respiratory function tests following standard guidelines. This minimised the risks associated with self-reported measures of health to which our analysis was vulnerable, such as differential reporting and recall bias.

However, our study also has limitations. Given this is a cross-sectional analysis, we have only been able to examine possible associations between coal mine fire smoke and asthma; we cannot infer causation. Also, we did not collect information on all potential confounders such as individual income, which may have influenced health service and medication usage, or medication adherence, which may have affected asthma severity and control. This analysis is nested within the Adult Survey cohort of the longitudinal Hazelwood Health Study, so follow up of these participants over time will provide further evidence regarding the impact of the Hazelwood mine fire event on Morwell asthmatics, and facilitate evaluation of additional confounders.

Our study may be subject to a number of other biases. We drew our sample of asthmatics from the Adult Survey participants, and by chance we may have sampled a greater proportion of mild asthmatics from Morwell compared to Sale. Furthermore, selection bias due to the volunteer effect and non-response bias may have impacted on whether the results are actually representative of the asthmatic populations in Morwell and Sale. It is difficult to say which way these may have biased our results.

Finally, our participants were drawn from a study population that had been determined by sample size calculations powered to detect a change in FEV_1_ over time for the Respiratory stream longitudinal cohort study. This may mean that our study was underpowered to detect small differences in asthma symptoms or lung function between the exposed and unexposed asthmatics. However, potential effect sizes between the two groups for the exposure were unknown and could not be calculated readily.

Three and a half years after the Hazelwood mine fire, our study has found some evidence of poorer asthma control in participants with asthma who were exposed to the coal mine fire smoke, but there was no significant difference in severity of asthma symptoms or lung function. Given the absence of literature, further research is needed in order to determine the clinical importance of our findings and whether coal mine smoke exposure has a causal role in long-term asthma control. The planned longitudinal study will further examine this question for both asthma control and severity. Investigating the association between individual PM_2.5_ exposure and asthma-related outcomes would help to characterise the degree of risk more comprehensively.

The chance of coal mine fires occurring in the future is likely to increase due to climate change[5], and this could result in increases in both the acute and chronic disease health burden. Evidence for the impact of coal mine fires smoke on health will allow governments and public health policy makers to plan for future similar events in order to minimise harm to populations at risk of exposure.

## Acknowledgments

The Respiratory Stream clinics were set up in facilities provided by the Central Gippsland Health Service, Sale, and The Healthcare Centre, Morwell. We thank Ms Susan Denny who oversaw all aspects of participant recruitment and Sharon Harrison for assistance with purchasing, logistics and set up of the clinic. Anthony Del Monaco is thanked for assistance with data management.

